# ACDA: Implementation of an Augmented Drug Synergy Prediction Algorithm

**DOI:** 10.1101/2022.10.21.513259

**Authors:** Sergii Domanskyi, Emily L. Jocoy, Anuj Srivastava, Carol J. Bult

## Abstract

**Motivation:** Drug synergy prediction is a complex problem typically approached with machine learning techniques using molecular data, pharmacological data, and knowledge of biological-interaction networks. The recently published Cancer Drug Atlas (CDA) uses a logistic regression model to predict a binary synergy outcome in cell-line models by utilizing drug target information, knowledge of genes mutated in each model, and the models’ monotherapy drug sensitivity. However, we observed low performance, 0.33, of the CDA measured by Pearson correlation of predicted versus measured sensitivity when we evaluated datasets from six studies that were not considered during the development of the CDA. Here we describe improvements to the CDA algorithm, the Augmented CDA, that improved performance by 71% and robustness to dataset variations in drug response values.

**Results:** We augmented the drug-synergy prediction-modeling approach CDA described in Narayan et al. by applying a random forest regression and optimization via cross-validation hyper-parameter tuning. We benchmarked the performance of our Augmented CDA (ACDA) compared to the original CDA algorithm using datasets from DrugComb, an open-access drug-combination screening data resource. The ACDA’s performance is 71% higher than that of the CDA when trained and validated on the same dataset spanning ten tissues. The ACDA performs marginally better (6% increase) than the CDA when trained on one dataset and validated on another dataset in 22 cases that cover seven tissues. We also compared the performance of ACDA to one of the winners of the DREAM Drug Combination Prediction Challenge (Mikhail Zaslavskiy’s algorithm which we denoted as EN). The performance of EN was smaller than that of the ACDA in 15 out of 19 cases. In addition to data from cell lines, we also trained the ACDA algorithm on Novartis Institutes for BioMedical Research PDX encyclopedia (NIBR PDXE) data and generated sensitivity predictions for the cases where drug-combination tumor-volume measurements were unavailable. Finally, we developed an approach to visualize synergy-prediction data using dendrograms and heatmaps instead of the Voronoi diagrams used in the CDA. The latter has a complex algorithmic realization and no publicly available implementation, whereas the ACDA visualization approach is more transparent and has open access. We implemented and wrapped the ACDA algorithm in an easy-to-use python package available from PyPI.

**Availability:** The source code is available at https://github.com/TheJacksonLaboratory/drug-synergy, and the software package can be installed directly from PyPI using pip.

## 1 Introduction

Drug combination therapy is a promising approach to cancer treatment (Mokhtari et al. 2017). Combination of cancer treatment with immunotherapy (Nam et al. 2019), immune checkpoint inhibitors (Patel & Minn 2018), and extension of drug repurposing (Sun et al. 2016) via combination therapy are widely researched. It is typical that a combination of drugs has no synergy (simultaneously targeting independent pathways) (Palmer & Sorger 2017). However, a more desirable approach is to use synergistic combinations that increase the therapeutic response rate and have, compared to monotherapies, lower potential to develop resistance to treatment (Gayvert et al. 2017).

Significant progress has been made in the development of computational methods for *in silico* testing of large number of compound combinations using existing high-throughput drug-screen data (Bansal et al. 2014; Menden et al. 2019; Torkamannia et al. 2022; Wu et al. 2022) to prioritize synergistic candidate pairs for further in *vitro* or in *vivo* testing. Moreover, efforts are ongoing to develop deep-learning models to identify synergistic combinations (reviewed in Baptista et al. 2022). However, to be robust, any such model will require a larger number of experimentally measured drug combinations for training than are currently available.

A recently developed method, Cancer Drug Atlas, CDA (Narayan et al. 2020) for drug-synergy prediction in cell-line models uses drugtarget information, knowledge of genes mutated in cell lines, and monotherapy drug sensitivity. Specifically, the authors developed an approach to calculate drug sensitivity-based drug-drug distance and further integrated knowledge of drug targets with models’ mutations. To substantiate use of drug-drug distance in the CDA approach, Narayan et al. performed an *in vitro* drug-synergy screen of 30 drug combinations in 9 glioblastoma (GBM) cell lines. Criteria for drug selection were that they have large drug-drug distance and high sensitivity in GBM cell lines. A significant number of the tested drug combinations showed synergy *in vitro*. In another set of experiments, using orthotopic mouse models with glioma (U87-MG-FM PDX model), triple-negative breast cancer (MDA-MD-231 -FM PDX model), melanoma (CHL1-FM PDX model), or leukemia (BV-173-Gluc PDX model), the authors of the CDA confirmed the validity and translational value of the CDA drug-synergy predictions.

The CDA was established on Genomics of Drug Sensitivity in Cancer (GDSC) data (Yang et al. 2013) and validated on a set of 437 manually curated tissue-specific drug pairs from the literature. These curated pairs are known to have synergy and span 240 drugs and 109 cell lines over several biological tissues. We implemented an in-house implementation of the CDA and re-evaluated it on the GDSC data, demonstrating results consistent with the Narayan et al. 2020 publication. However, we observed low performance of the CDA when trained and tested on several datasets from DrugComb (Zheng et al. 2021), detailed below in the Results section.

Here, we describe the implementation of improvements to the CDA drug-synergy prediction modeling approach by applying a Classification And Regression Tree (CART)-based model instead of linear regression. Further, we added training optimization via cross-validation hyperparameter tuning and implemented the algorithm as a Python package called Augmented CDA (ACDA). We benchmarked the CDA and ACDA methods on a collection of datasets. Additionally, to support the conclusions of ACDA strengths, we implemented and tested the ACDA alongside a second-best method developed by Mikhail Zaslavskiy (which we denote as EN) of the community-powered DREAM Drug Combination Prediction Challenge to develop and assess computational models for drug-synergy prediction (Menden et al. 2019). We did not use the best method of the DREAM Drug Combination Prediction Challenge, as it does not satisfy the ACDA data requirements. Additionally, we trained and benchmarked ACDA predictions of therapeutic response in Patient-derived Xenograft (PDX) models from the Novartis Institutes for BioMedical Research PDX encyclopedia (NIBR PDXE collection (Gao et al. 2015) encompassing 281 PDX models of several cancer types. As an example of ACDA application in prioritizing candidate drug combinations for testing in PDX models, we generated a set of model drug-drug candidates for the NIBR PDXE breast cancer subset where drug-combination tumor-volume measurements were unavailable. Finally, we simplified the visualization of drug-sensitivity clustering dendrograms on a two-dimensional layout by using dendrograms and heatmaps in place of the Voronoi diagrams used in the CDA.

## 2 Methods

### 2.1 ACDA algorithm description

The Augmented CDA model followed similar steps to those employed by the CDA model (Narayan et al. 2020). Step 1 is the same as in the CDA, and Steps 2-4 are modified compared to the CDA:

Step 1. To calculate drug-drug distance (a drug-similarity measure based on monotherapy sensitivity in the models), first project drug response vectors (n drugs, m cell lines) sensitivity measures (e.g., AUC) to cosine similarity, and construct matrix A. Then calculate Euclidean similarity M (n by n drugs) of A, hierarchically cluster M with Ward linkage to make a dendrogram. Finally, calculate the cophenetic distance of the clustering.
Step 2. Build a random forest regression model using the monotherapy drug sensitivities, target information, and the drug-drug distance.
Step 3. Train the regression model and compute (predict) synergy values for new or validation data.
Step 4. Visualize predicted and known synergy-pairs data using dendrograms and heatmaps (compare to Voronoi diagrams used in the CDA).

The ACDA has the following novel aspects compared to CDA: in Step 2 random forest regression is used instead of logistic regression, and random search is used for hyperparameter tuning via k-fold cross-validation. In Step 2, compared to a linear model fit evaluation done by Narayan et al. 2020, we perform cross-validation using Monte Carlo Cross-Validation approach according to which a randomly selected 2/3 portion of data was used for training, then the remaining 1/3 of the data was used for model performance evaluation (Kuhn & Johnson 2013 p.). The random splitting, training and evaluation was repeated 10 times to benchmark the ACDA and CDA methods. Step 4 implementation is detailed in Section 2.3.

The DrugComb dataset (Zheng et al. 2021) contains all measured drug pairs regardless of the synergy value, whereas in GDSC drug-synergy pairs have binary “synergy, no synergy” values; therefore, we applied the CDA with linear regression instead of logistic regression.

Since the ACDA is based on several features from different data types (pharmacology and molecular), data curation is a crucial step. The underlying software uses models, drugs, and gene identifiers to match and query the data tables. These identifiers were curated so that they were consistent across the tables. Curated data is shared for reproducibility, and documentation is available at https://acda.readthedocs.io.

Hyperparameter optimization was performed by a randomized search with k-fold cross-validation (k=3) using one data subset as a training dataset. The best estimator was determined by performance on another subset.

### 2.2 Data Description

We used pharmacology and molecular data from (Yang et al. 2013), including GDSC drug sensitivity, model mutations, and drug-targets data (Supplementary Table 1). We manually curated the CDA synergy pairs (Narayan et al. 2020) to harmonize drugs and model names. The GDSC set contains subsets GDSC1 and GDSC2, which are older and newer versions of GDSC, respectively. Notably, the GDSC2 subset has been screened using an improved GDSC1 screen design and assay (Yang et al. 2013).

We tested ACDA vs. CDA on breast cell lines of various cancer types from the GDSC1 and GDSC2 subsets (the two editions of the GDSC drug screens) with CDA synergy pairs. The GDSC1 breast subset contains 119 drug pairs (54 with synergy and randomly chosen 65 unknown pairs assuming no synergy). The GDSC2 breast subset contains 163 pairs (74 with synergy, 89 without synergy), and GDSC1&2 with 191 pairs (112 with synergy, 79 without synergy). For each scenario, we tested 10 randomly chosen sets of “no synergy” pairs.

The second data source was DrugComb which includes 650,909 harmonized drug sensitivity and synergy values from many *in vitro* studies and tissues (Zheng et al. 2021)(see Supplementary Figures 1 and 2) and, specifically, the AstraZeneca dataset from Menden et al. 2019. The data in DrugComb do not have model mutation information; therefore, where available, we used GDSC mutations for DrugComb. We considered only those DrugComb sets for which there are at least 1,000 drug-drug-model entries per study per tissue. Because the DrugComb-AstraZeneca dataset is much larger than the GDSC known synergy set, we use the DrugComb subset in a down-sampling experiment by using a reduced number of drug-drug-model entries in a training set and a constant number of drug-drug-model entries in the validation set.

As an independent dataset for demonstrating high benchmarking performance and utility of the ACDA to generate candidate drug-drug combinations in PDX models, we use the NIBR PDXE collection (Gao et al. 2015). The dataset contains tumor-volume response data of 281 models of six cancer types treated with single drugs and drug combinations: colorectal cancer (CRC), gastric cancer (GC), pancreatic ductal adenocarcinoma (PDAC), breast cancer (BRCA), non-small cell lung cancer (NSCLC), and cutaneous melanoma (CM).

### 2.3 Visualization

There are two visualization tools in the ACDA. The first tool allows clustering of monotherapy sensitivities into a dendrogram of drug-drug similarity and depicts the known synergy pairs as arcs ordered according to the order of the dendrogram leaves. Lighter colors denote a smaller cophenetic distance between drugs, and darker colors show that drug-drug cophenetic distance is large. The dendrogram is split into 10 clusters. This dendrogram representation allows for visually assessing whether known synergy pairs tend to have large cophenetic distances, as was discussed in the work of Narayan et al. 2020. The second tool presents predicted drug-synergy pairs as a clustered heatmap where each point on the heatmap corresponds to a drug pair. Dendrograms reflect clustering of similarity of drug sensitivities.

### 2.4 Application to PDX models

To analyze PDX model data, we used sensitivity to monotherapy (63 drugs) and tumor mutation profiles of the PDX models from the NIBR PDXE collection (Gao et al. 2015). First, we performed benchmarking via Monte Carlo Cross-Validation Scheme repeated ten times for each of the four methods (ACDA, CDA, EN, and EN-ACDA) and each of the six cancer types, listed in Section 2.2, in NIBR PDXE (see Supplementary Methods for methodological details). Next, we used all available drug combinations in the breast cancer (BRCA) subset of NIBR PDXE to train the methods and then generated sensitivity predictions for the cases where drug-combination tumor-volume measurements were unavailable.

## 3 Results and Discussion

### 3.1 Performance of the ACDA synergy prediction algorithm

The average Pearson correlation coefficient of predicted versus measured drug-combination sensitivity values and SEM (standard error of the mean) for each case of the GDSC and DrugComb-AstraZeneca datasets are reported in Table 1. In the case of the GDSC1-breast dataset, the performance of the ACDA was consistently higher or nearly equal to that of the CDA. Notably, the CDA correlation coefficient values were similar to those in the CDA publication results (Narayan et al. 2020). However, we showed that the choice of a logistic model had a negative effect on the accuracy of drug-response prediction when using the DrugComb-AstraZeneca drug screening dataset (and several other GDSC subsets, Table 1). By constructing a CART-based model for the regression and classification problem, we overcame the assumptions necessary for using a logistic model and improve the quality of model performance.

**Table 1.** Benchmark Pearson correlation coefficient of ACDA/CDA models on select tissues and studies.

| Dataset | Tissue | Train | Test | ACDA | CDA |
| --- | --- | --- | --- | --- | --- |
| GDSC1&2 | Breast | 127 | 64 | $0.572 \pm 0.029$ | $0.468 \pm 0.032$ |
| GDSC1 | Breast | 79 | 40 | $0.617 \pm 0.044$ | $0.620 \pm 0.038^{**}$ |
| GDSC2 | Breast | 108 | 55 | $0.443 \pm 0.044$ | $0.306 \pm 0.045$ |
| GDSC1&2 | Bladder | 73 | 37 | $0.518 \pm 0.045$ | $0.461 \pm 0.034$ |
| GDSC1 | Bladder | 60 | 30 | $0.686 \pm 0.051$ | $0.624 \pm 0.051$ |
| GDSC2 | Bladder | 58 | 29 | $0.667 \pm 0.046$ | $0.427 \pm 0.026$ |
| DC-AZ | Breast | 6585 | 3293 | $0.885 \pm 0.016$ | $0.57 \pm 0.017^*$ |
| DC-AZ | Lung | 2340 | 1170 | $0.539 \pm 0.016$ | $0.335 \pm 0.015^*$ |
| DC-AZ | Urinary tract | 2921 | 1461 | $0.760 \pm 0.11$ | $0.505 \pm 0.008^*$ |

\*Linear Regression was used instead of Logistic Regression, \*\*Similar performance of CDA compared to ACDA is observed, DC-AZ is DrugComb-AstraZeneca.

We observed that the performance of the ACDA on a down sampled DrugComb-AstraZeneca dataset decreased slightly with the smaller training set. In contrast, the performance of the CDA is much lower on the same set. However, the performance of the CDA did not significantly change with the training set size (Supplementary Figure 3).

We performed additional validation and benchmarking of our ACDA and CDA implementation using datasets from DrugComb (Zheng et al. 2021). We compared our model performance to that of one of the top-performing models, EN, from the DREAM challenge for drug-synergy prediction (Menden et al. 2019). When trained and tested on the same dataset, the EN method shows a performance of 0.442±0.032, which is comparable to those of the ACDA (0.57±0.04) and the CDA (0.334±0.049). However, EN performs poorly (0.009±0.012) when training and testing are done on different studies. In the latter scenario, the ACDA and the CDA have similar but low performances: 0.228±0.046 and 0.215±0.051, respectively. A combination of EN and ACDA is a viable approach (Supplementary Figures 4 and 5), which aligns the strengths of both methods in a generalizable scenario giving 0.645±0.036, or a 93% increase compared to the CDA when trained and tested on subsets of the same study.

The hyperparameter tuning by training on the GDSC2 breast subset and testing on the GDSC1 breast subset led to the increased correlation of the predicted values with ground truth from 0.390 (base model) to 0.797. A similar test using GDSC1 and GDSC2 as a training and testing set suffered from overfitting and a decrease in correlation from 0.248 (base model) to 0.209.

### 3.2 Visualization of the synergistic pairs

The visualization example of clustering drug-sensitivity measure values overlaid with known drug-synergy values for the DrugComb-Astra-Zeneca breast cancer subset is shown in Supplementary Figure 6. This dendrogram representation with color-coded arcs allows researchers to visually determine that known synergy pairs tend to have large co-phenetic distances. Another visualization example is a heatmap of the synergy-pairs predictions for the GDSC2 breast subset, where each point on the heatmap corresponds to a drug pair. The dendrograms reflect clustering of similarity of drug sensitivities (Supplementary Figure 7).

### 3.3 Application to PDX models

Using Monte Carlo Cross-Validation we applied the ACDA method trained on a randomly selected 2/3 portion of measured dual-drug responses to predict synergistic combinations (model-specific drug pairs) in NIBR PDXE models (Gao et al. 2015), for which tumor-response values were held out from the training set. The data splitting, training and model evaluation was repeated 10 times. ACDA, CDA, EN, and EN-ACDA results for each of the six cancer types in NIBR PDXE are detailed in Supplementary Figures 8 and 9. ACDA, CDA, and EN-ACDA had similar performances, while EN showed significantly lower performance with GC and PDAC subsets.

Then we used all available drug-combinations data in the breast cancer subset of NIBR PDXE to train the methods and then generated sensitivity predictions for the cases where drug-combination tumor-volume measurements were unavailable, but the computational model predictors were available. The top 20 drug-drug-model synergy candidates are listed in Supplementary Table 2, with model X-4567 appearing in the top 20. A complete list of predicted sensitivity to drug combinations predicted from ACDA, CDA, EN, and EN-ACDA for each model is provided in Supplementary Table 3.

## 4 Conclusion

The newly developed ACDA method is available as an open-source software package and includes the CDA, EN, and EN-ACDA implementations. Benchmarking the four methods demonstrated that ACDA and EN-ACDA were better in predicting synergistic drug combinations compared to CDA and EN. The ACDA approach can be used with cell-line drug screens and *in vivo* PDX model tumor-volume data to predict drug-drug combinations that may synergistically affect specific models.

The current ACDA synergy predictions are partly dependent on the experimental system model gene mutations. Future versions of ACDA will include genes downstream of the drug targets using known protein-protein interaction networks known as parsimonious composite networks (Huang et al. 2018). Expanding the gene lists to include nearest neighbor genes based on protein-protein interaction networks will also be evaluated to see if the performance of ACDA can be enhanced. Finally, future studies will integrate into ACDA information on the overlap of mutations with sets of ontology pathways to define a new covariate in the regression. For example, PDX model X-4567 is predicted to be sensitive to paclitaxel and binimetinib (Supplementary Table 2). This model has mutations in nine of the 267 MAPK pathway related genes (Subramanian et al. 2005). The combination of paclitaxel and binimetinib was studied in clinical trial NCT01649336 and was shown to have clinical benefit in treating epithelial ovarian tumors harboring alterations in the MAPK pathway (Grisham et al. 2018). Thus, adding pathway context to the model may further enhance the drug synergy predictions.

## Acknowledgements

We thank Brian J. Sanderson for helpful discussions and feedback and for evaluating the ACDA algorithm. We also thank Grace A. Stafford for CDA synergy data curation.

## Data availability

All the data collections including the DrugComb, GDSC, and NIBR PDXE datasets are publicly available. The ACDA source code, documentation, examples of usage, and example data are openly available at https://github.com/TheJacksonLaboratory/drug-synergy and https://acda.readthedocs.io/. The list of curated CDA synergy pairs can be found in Supplementary Table 5.

## Supplementary Data

### 5 Supplementary Figures

**Figure S1.**
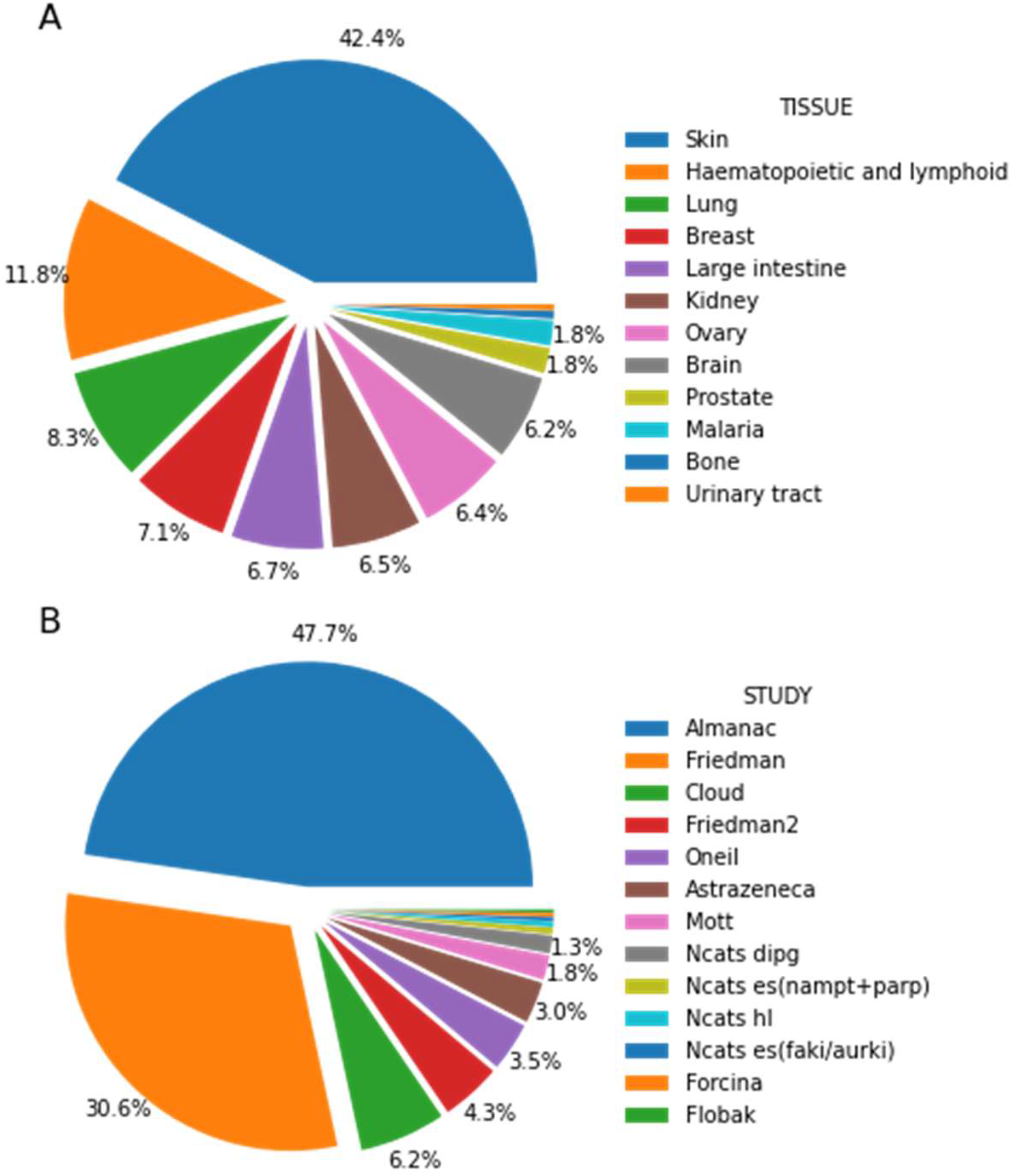
DrugComb data composition overview by tissues and by studies. Distribution of drug pairs in DrugComb (Zheng et al. 2021), a comprehensive drug-sensitivity data repository and analysis portal, separated by (A) studies and (B) tissues. Only those study-tissue cases with at least 1,000 measured drug pairs are shown. Overall, DrugComb collects more than 6500,00 model-drug-drug entries, spanning 26 studies, 288 cell-line models, and 4,268 drugs. Skin is the best-represented tissue in this data resource, accounting for 42.4% of all data entries. Similarly, the “Almanac” and “Friedman” studies account for 78.3% of the data; however, other valuable studies such as “AstraZeneca” and “Oneil” are included as well.

**Figure S2.**
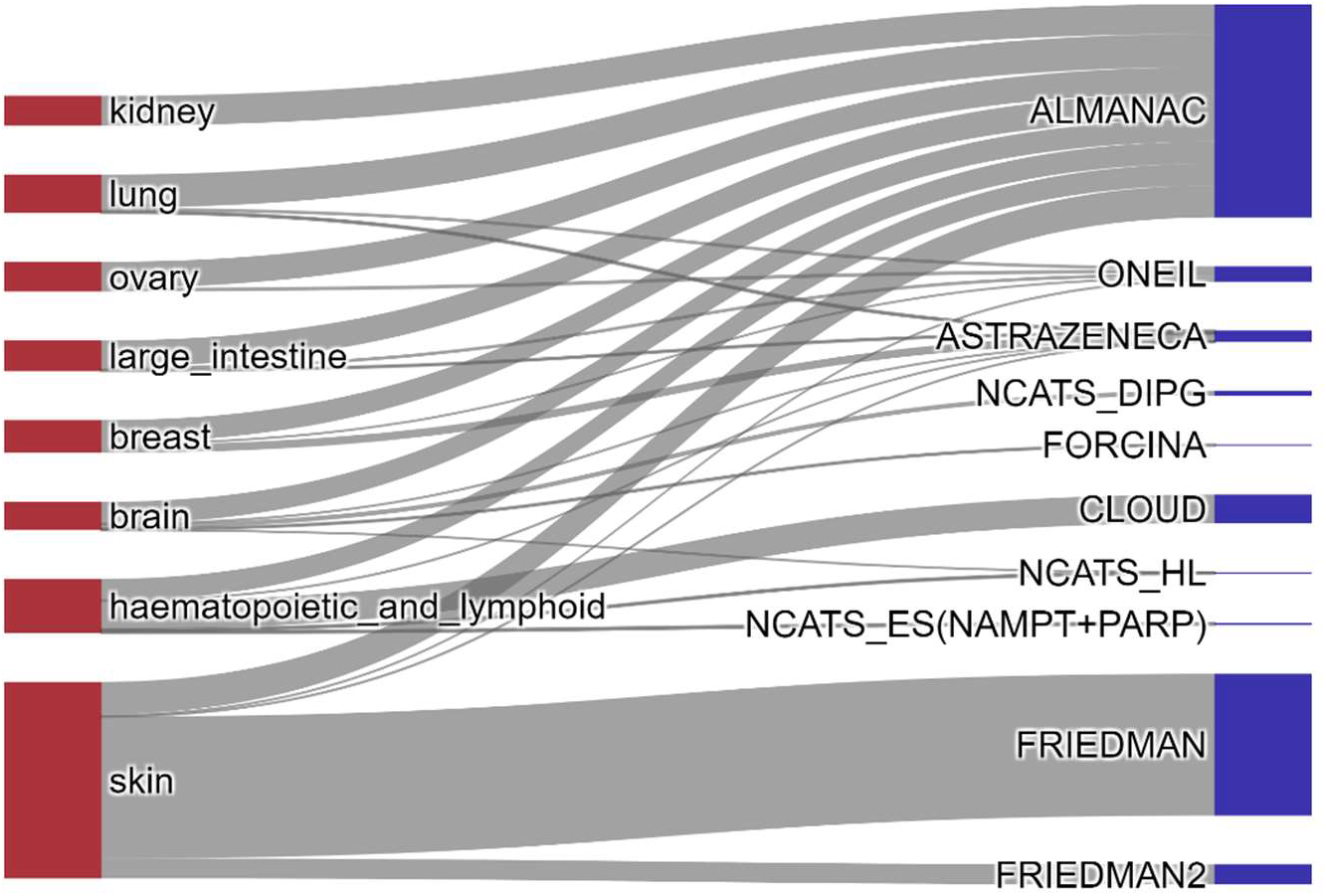
Comparison of the tissues and studies from DrugComb highlighting weight of each study by the data quantity. DrugComb collects measured drug pairs from several tissues and studies. Only those study-tissue cases that have at least 1,000 measured drug pairs are shown. Noticeably, a large fraction of all pairs is from skin. The largest studies in this dataset are “Almanac” and “Friedman” (Zheng et al. 2021). The figure was generated with the visualization tools of Digital Cell Sorter (Domanskyi et al. 2019, 2021).

**Figure S3.**
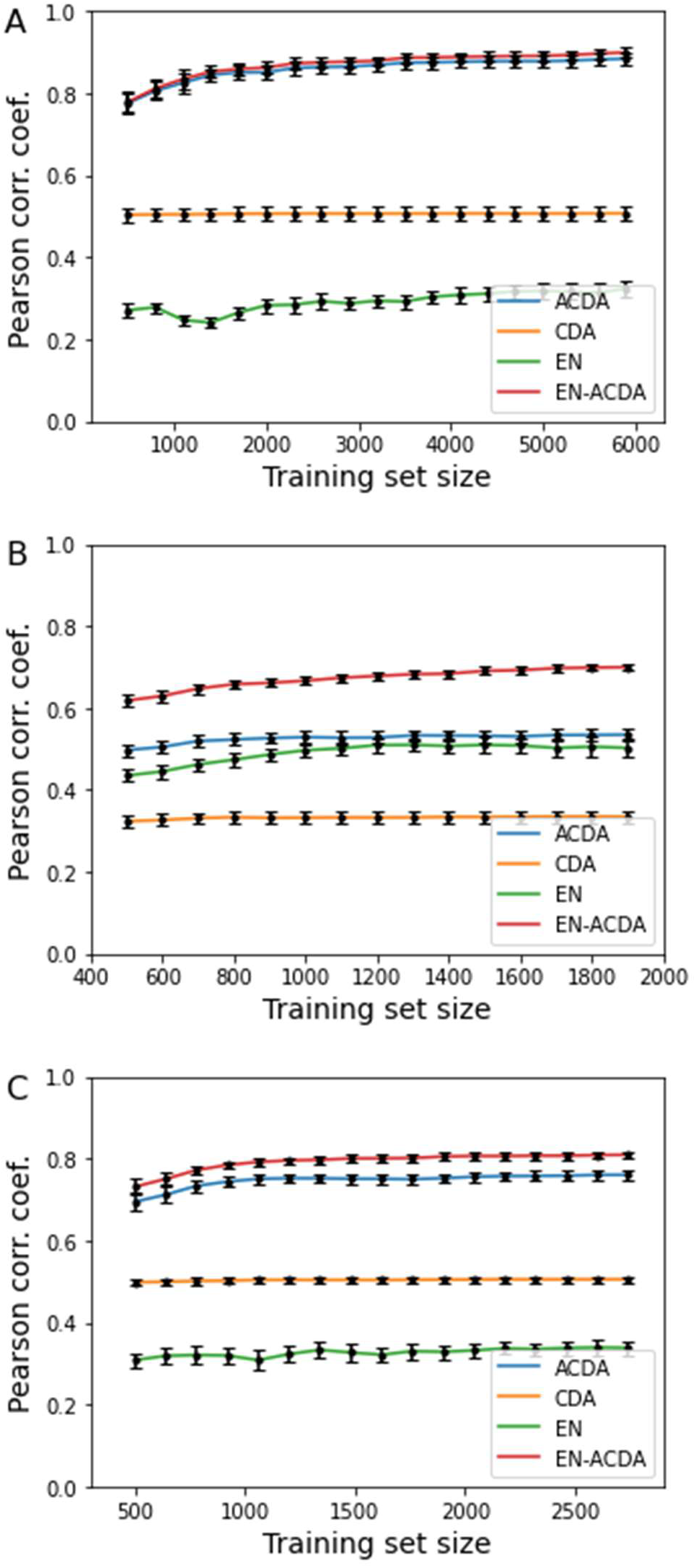
Effect of down-sampling for DrugComb AstraZeneca training dataset. The panels correspond to three tissues from the AstraZeneca dataset and contain (A) 9,878 entries in the breast, (B) 3,510 entries in the lung, and (C) 4,382 entries in the urinary tract set. Training is done on a subsample of 2/3 of the randomly split data, and testing is done on 1/3 of the data. For each training set size, random split and subsampling are repeated 10 times. The random splits are reused across all training set sizes. A consistent effect is observed: (i) ACDA performance is higher than that of CDA, while both ACDA/CDA have nearly constant performance across a range of training set sizes, (ii) EN performance is significantly lower than that of the other methods in (A) and (C), (iii) Addition of EN features to ACDA (see EN-ACDA labelled curves) improves ACDA performance.

**Figure S4.**
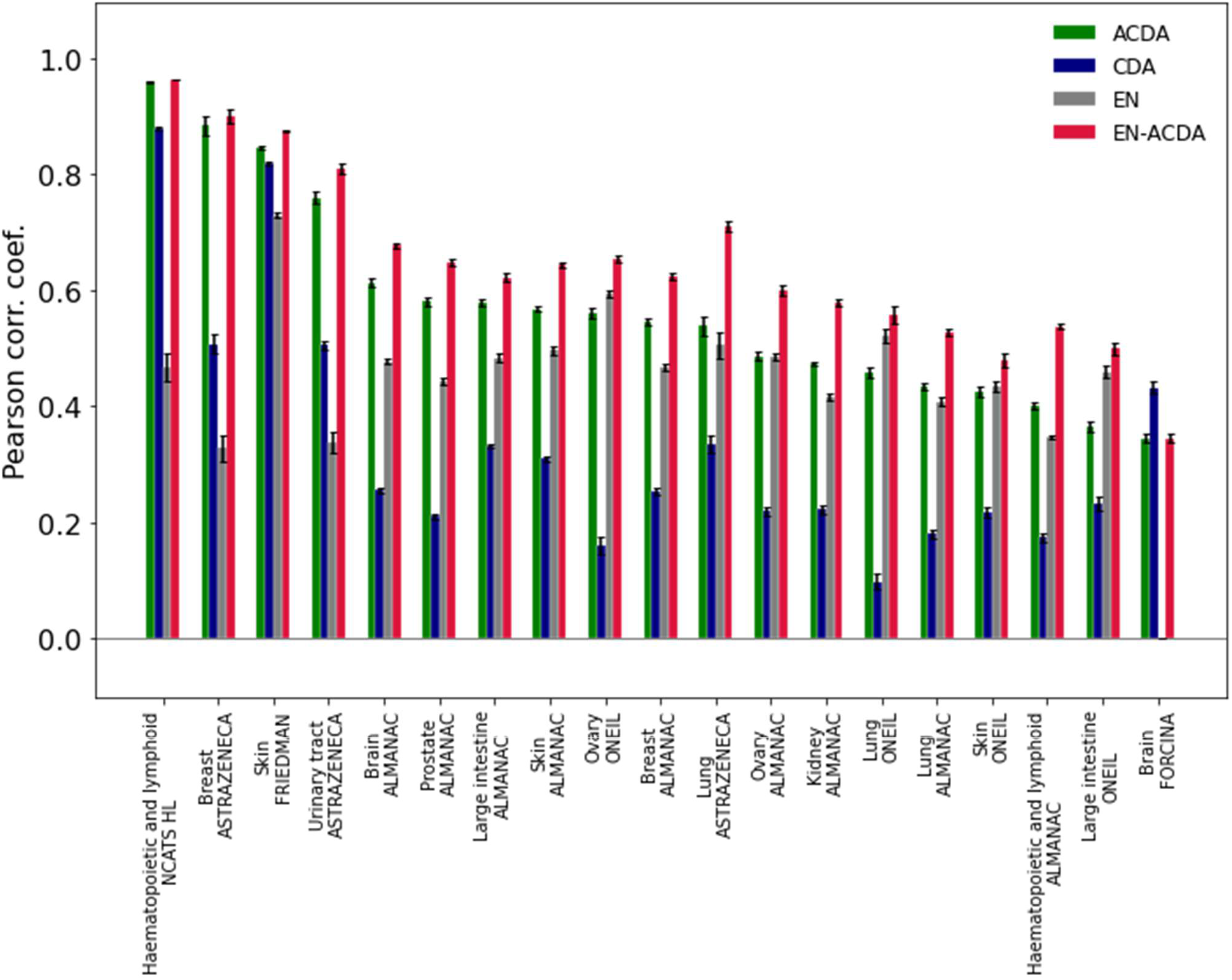
Comparison of drug-synergy prediction methods across a range of datasets from DrugComb by training and testing on data within the same studies. Benchmarking the synergy-prediction methods on DrugComb datasets for which there are at least 1,000 entries per tissue and study, after any entries with missing values were removed. Model mutations data are taken from the GDSC Sanger dataset where available. Training is done on 2/3 of the data and testing on 1/3 of the data of the same dataset, repeated over 10 random splits in a Monte Carlo Cross-Validation scheme. The chart shows Pearson correlation coefficients of the experimentally measured synergy values with values predicted by the four methods. Black error bars show SEM over 10 Monte Carlo iterations. ACDA performance is consistently higher than that of CDA for most of the tested datasets. Use of EN features with a random forest regression (EN-labelled curves) leads to the highest performance for most of the datasets. The EN-ACDA combination shows the highest performance across nearly all tested datasets.

**Figure S5.**
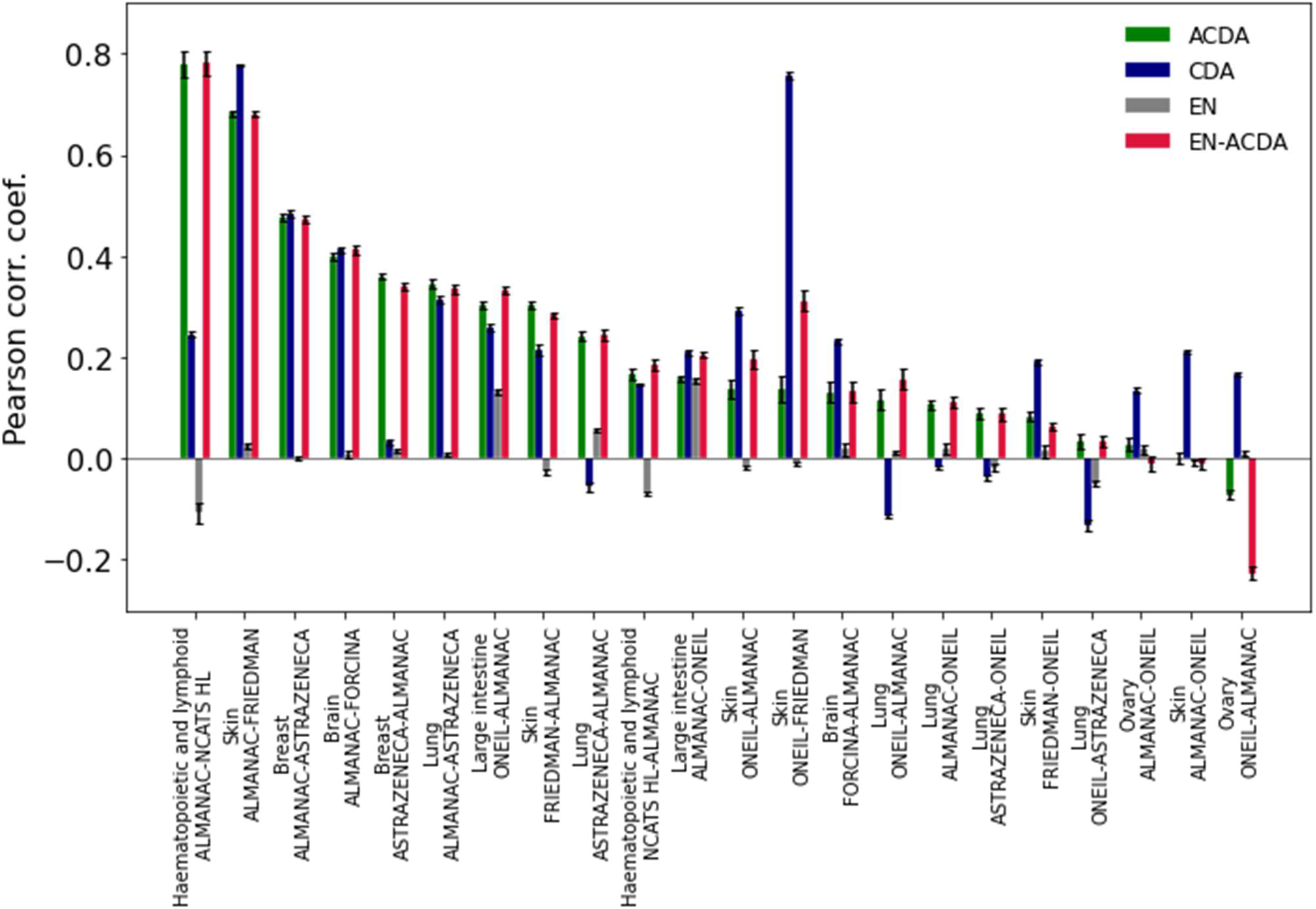
Comparison of drug-synergy prediction methods across a range of datasets from DrugComb by training and testing on data from one study and testing on a different study. Benchmarking synergy prediction methods on DrugComb datasets for which there are at least 1,000 entries per tissue and study, after any entries with missing values were removed. Note that for model mutations, data is taken from GDSC Sanger dataset where available. Training is done on a 1/2 of data from one study, testing on 1/2 of data of another study specified in the figure labels, repeated over 10 random splits in a Monte Carlo Cross-Validation-like scheme. The chart shows Pearson correlation coefficients of the experimentally measured synergy values with values predicted by the four methods. Black error bars show SEM over 10 Monte Carlo iterations. ACDA, CDA and EN-ACDA have similar performance. Because here the datasets share limited overlap, the EN approach alone gives near zero correlation with ground truth values; therefore, it cannot be used in cases when training and testing data do not overlap in models and drugs.

**Figure S6.**
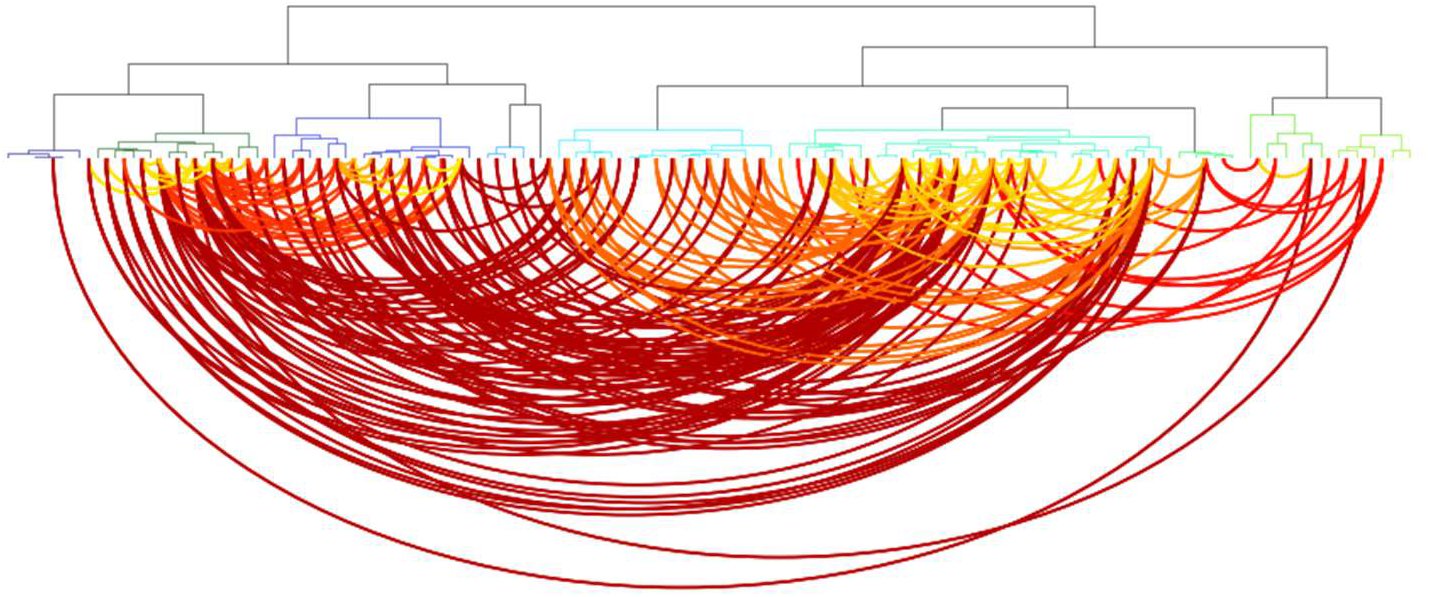
Clustering of drug-sensitivity measure values overlaid with known drug-synergy values. DrugComb AstraZeneca breast cancer cell-line sensitivity (AUC) is clustered into a dendrogram of drug-drug similarity (top panel), and visualization of the known synergy pairs (bottom panel) is ordered according to the order of the dendrogram leaves. The pairs with synergy score at least 20 are assumed to be known synergy pairs. Lighter colors (yellow) denote smaller cophenetic distance between drugs, darker colors show that drug-drug cophenetic distance is large. The dendrogram is split into 10 clusters and color-coded. This dendrogram representation allows visual assessment of whether known synergy pairs tend to have large cophenetic distances, as was discussed in the work of (Narayan et al. 2020).

**Figure S7.**
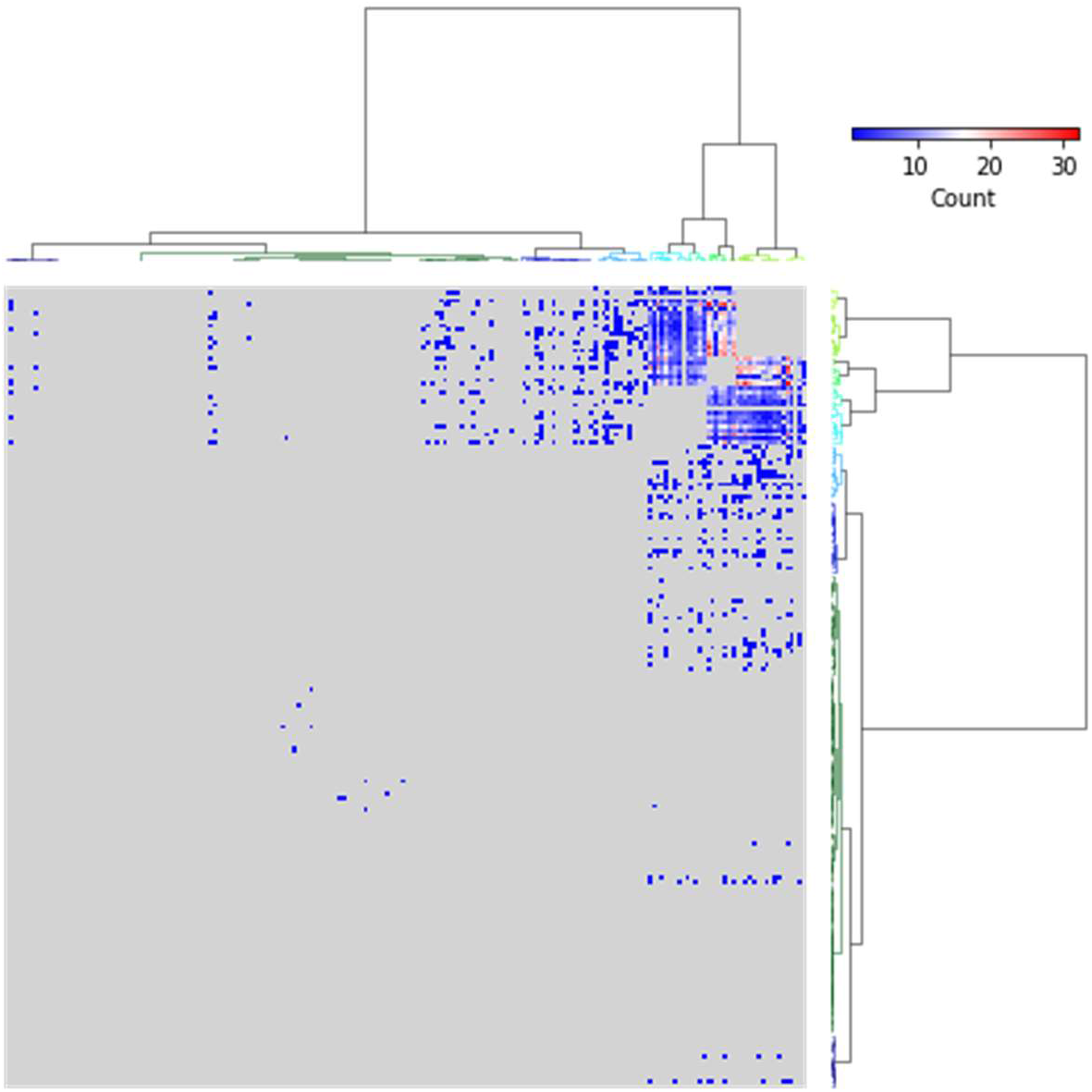
Heatmap of the synergy-pairs predictions for GDSC2 breast subset. Each point on the heatmap corresponds to a drug pair. Dendrograms reflect clustering of similarity of drug sensitivities. The color reflects number of cell-line models in which the drug combination may have a synergistic effect, with regression values at least 0.95. We observe that the top right area of the heatmap tends to contain many synergistic combinations, i.e., a small subset of drugs from the top right section of the dendrogram is predicted to have many synergistic pairs among them.

**Figure S8.**
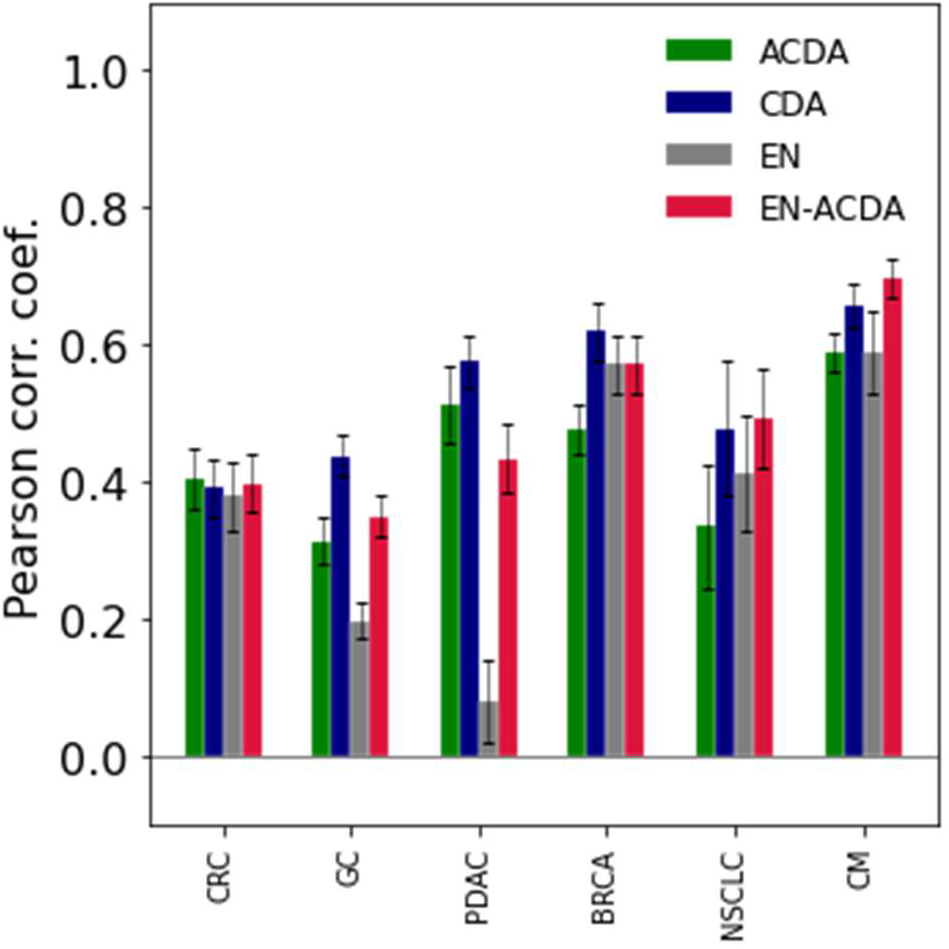
Comparison of drug-synergy prediction methods across a range of cancer types from NIBR PDXE dataset. Benchmarking synergy prediction methods on PDX models of six DrugComb datasets (one dataset for each of six cancer types) for which monotherapy and mutation profiles are available. There are at least 117 drug-drug model entries with non-missing features for each cancer type (CRC: 294, GC: 270, PDAC: 219, BRCA: 188, NSCLC: 148, CM: 117). For each cancer type, training is done on 2/3 of the data, and testing on 1/3 of the data from the same cancer type, repeated over 10 random splits in a Monte Carlo Cross-Validation-like scheme. The chart shows Pearson correlation coefficients of the experimentally measured synergy values with values predicted by the four methods. Black error bars show SEM over 10 Monte Carlo iterations. ACDA, CDA and EN-ACDA have similar performances, while EN shows significantly lower performance with the GC and PDAC subsets.

**Figure S9.**
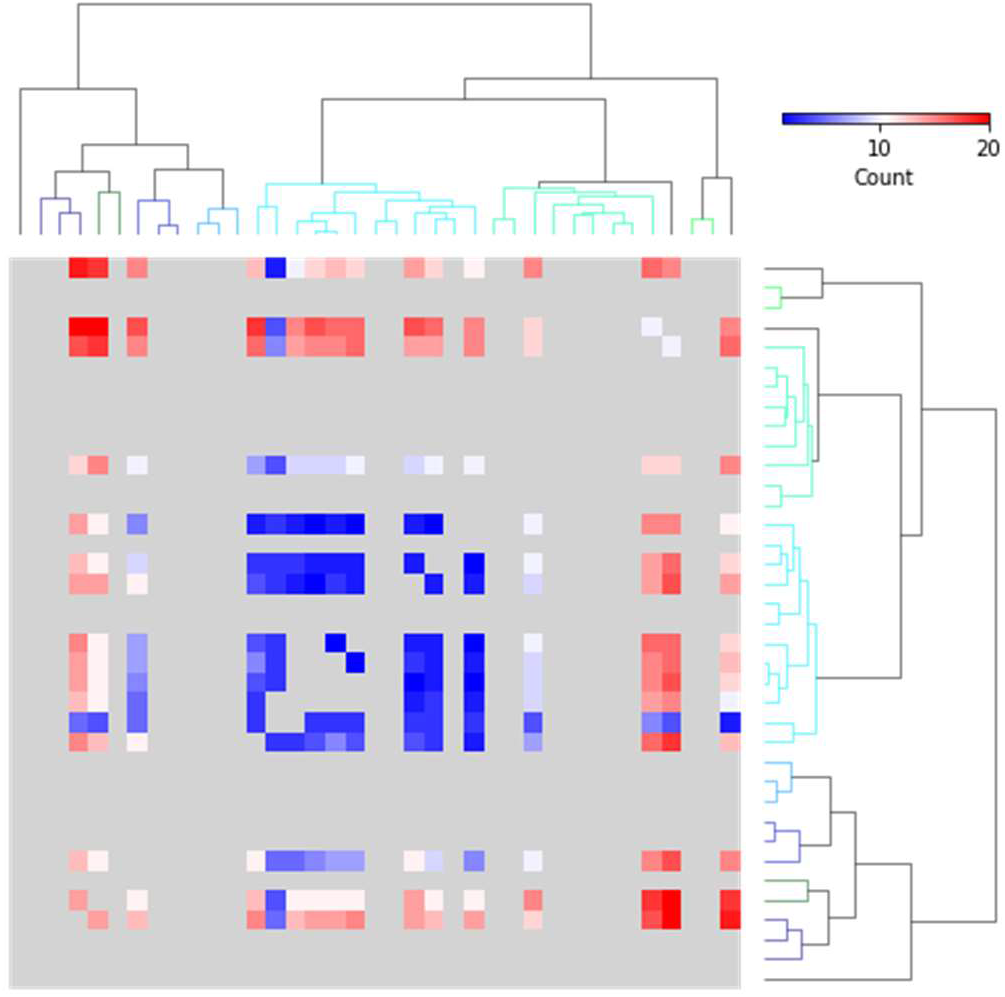
Heatmap of the synergy-pairs predictions by EN-ACDA for NIBR PDXE BRCA subset. Each point on the heatmap corresponds to a drug pair. Dendrograms reflect clustering of similarity of drug sensitivities. The color reflects the number of cell-line models in which the drug combination may have a synergistic effect, with predicted scores of at least 0. We observe that a small subset of drugs from the top right section of the dendrogram is predicted to have a synergistic effect in many of the models. Specifically, these drug combinations are binimetinib-paclitaxel, binimetinib-LLM871, BKM120-paclitaxel, BKM120-LLM871, CLR457-paclitaxel, and CLR457-LLM871.

### 6 Supplementary Tables

**Supplementary Table 1.**
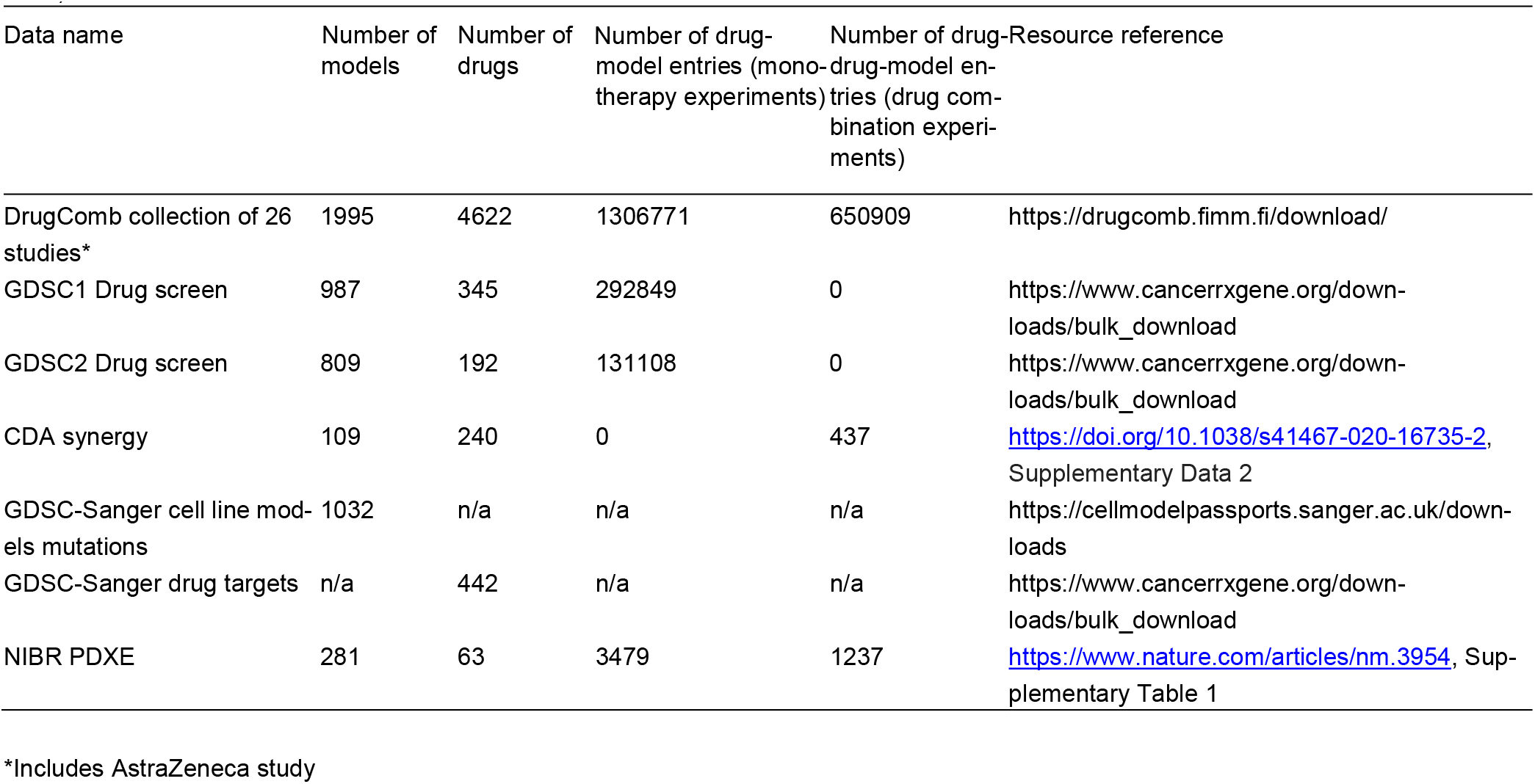
Description of the datasets used in this work. The sources of the data used in our work are DrugComb (Zheng et al. 2021), GDSC (Yang et al. 2013), and CDA (Narayan et al. 2020). The PDX resource used in our analysis is NIBR PDXE (Gao et al. 2015).

**Supplementary Table 2.** Top drug-synergy combination candidates among 39 breast cancer PDX models and 119 unique drugdrug combinations sorted by EN-ACDA score. By using EN-ACDA we train a classifier on PDX models for measured drug combinations and predict synergy for PDX models where drug-combination response is unknown. The top 20 synergy-pairs candidates are listed. A complete list of 9184 predicted sensitivity to drug-drug combinations for each model is shown in Supplementary Table 3.

| Model | Drug 1 | Drug 2 | ACDA | CDA | EN | EN-ACDA |
| --- | --- | --- | --- | --- | --- | --- |
| X-4567 | CLR457 | LJM716 | 0.724 | 0.845 | 0.849 | 0.821 |
| X-4567 | paclitaxel | LJM716 | 0.724 | 0.844 | 0.849 | 0.821 |
| X-4567 | BKM120 | LJM716 | 0.724 | 0.845 | 0.849 | 0.821 |
| X-4567 | LLM871 | LJM716 | 0.724 | 0.844 | 0.849 | 0.821 |
| X-4567 | LLM871 | BGJ398 | 0.692 | 0.422 | 0.843 | 0.818 |
| X-4567 | paclitaxel | BGJ398 | 0.692 | 0.422 | 0.843 | 0.818 |
| X-4567 | CLR457 | paclitaxel | 0.866 | 2.114 | 0.843 | 0.816 |
| X-4567 | CLR457 | LLM871 | 0.866 | 2.114 | 0.843 | 0.816 |
| X-4567 | BKM120 | LLM871 | 0.866 | 2.114 | 0.843 | 0.816 |
| X-4567 | BYL719 | LLM871 | 0.866 | 2.114 | 0.843 | 0.816 |
| X-4567 | BYL719 | binimetinib | 0.866 | 1.691 | 0.843 | 0.816 |
| X-4567 | BYL719 | paclitaxel | 0.866 | 2.114 | 0.843 | 0.816 |
| X-4567 | BKM120 | paclitaxel | 0.866 | 2.114 | 0.843 | 0.816 |
| X-4567 | CLR457 | binimetinib | 0.866 | 1.691 | 0.843 | 0.816 |
| X-4567 | LLM871 | CLR457 | 0.866 | 2.114 | 0.843 | 0.816 |
| X-4567 | paclitaxel | CLR457 | 0.866 | 2.114 | 0.843 | 0.816 |
| X-4567 | paclitaxel | BYL719 | 0.866 | 2.114 | 0.843 | 0.816 |
| X-4567 | paclitaxel | BKM120 | 0.866 | 2.114 | 0.843 | 0.816 |
| X-4567 | LLM871 | binimetinib | 0.866 | 1.691 | 0.843 | 0.816 |
| X-4567 | paclitaxel | binimetinib | 0.866 | 1.691 | 0.843 | 0.816 |

### 7 Supplementary Methods

In order to apply ACDA to PDX data from NIBR PDXE, the steps outlined below must be carried out:

1. Select the PDX models of one cancer type (e.g., BRCA).
2. Generate ACDA-formatted data by following instructions at https://acda.readthedocs.io/en/latest/data.html#alternative-data-format-simple and https://acda.readthedocs.io/en/latest/examples.html#examples.
3. Map the tumor volume-derived response category to numeric values, i.e., CR: 1, PR: 0.5, SD: −0.25, PD: −1, SD->PD: −0.75, SD-->-->PD: −0.5, CR-->PD: 0.75, CR-->-->PD: 0.75, PR-->PD: 0.25, PR-->-->PD: 0.25.
4. Train classifier on drug-drug model entries with no missing features for which response in known.
5. Generate response predictions on drug-drug-model entries with no missing features for which response is not known.

## References

Bansal, M., Yang, J., Karan, C., Menden, M. P., Costello, J. C., Tang, H., Xiao, G., et al. (2014). ‘A community computational challenge to predict the activity of pairs of compounds’, Nature Biotechnology, 32/12: 1213–22.

Baptista, D., Ferreira, P. G., & Rocha, M. (2022). ‘A systematic evaluation of deep learning methods for the prediction of drug synergy in cancer’. bioRxiv.

Gao, H., Korn, J. M., Ferretti, S., Monahan, J. E., Wang, Y., Singh, M., Zhang, C., et al. (2015). ‘High-throughput screening using patient-derived tumor xenografts to predict clinical trial drug response’, Nature Medicine, 21/11: 1318–25. Nature Publishing Group.

Gayvert, K. M., Aly, O., Platt, J., Bosenberg, M. W., Stern, D. F., & Elemento, O. (2017). ‘A Computational Approach for Identifying Synergistic Drug Combinations’, PLOS Computational Biology, 13/1: e1005308. Public Library of Science.

Grisham, R. N., Moore, K. N., Gordon, M. S., Harb, W., Cody, G., Halpenny, D. F., Makker, V., et al. (2018). ‘Phase Ib Study of Binimetinib with Paclitaxel in Patients with Platinum-Resistant Ovarian Cancer: Final Results, Potential Biomarkers, and Extreme Responders’, Clinical cancer research: an official journal of the American Association for Cancer Research, 24/22: 5525–33.

Huang, J. K., Carlin, D. E., Yu, M. K., Zhang, W., Kreisberg, J. F., Tamayo, P., & Ideker, T. (2018). ‘Systematic Evaluation of Molecular Networks for Discovery of Disease Genes’, Cell Systems, 6/4: 484–495.e5. Elsevier.

Kuhn, M., & Johnson, K. (2013). ‘Over-Fitting and Model Tuning’. Kuhn M. & Johnson K. (eds) Applied Predictive Modeling, pp. 61–92. Springer: New York, NY.

Menden, M. P., Wang, D., Mason, M. J., Szalai, B., Bulusu, K. C., Guan, Y., Yu, T., et al. (2019). ‘Community assessment to advance computational prediction of cancer drug combinations in a pharmacogenomic screen’, Nature Communications, 10/1: 2674. Nature Publishing Group.

Mokhtari, R. B., Homayouni, T. S., Baluch, N., Morgatskaya, E., Kumar, S., Das, B., & Yeger, H. (2017). ‘Combination therapy in combating cancer’, Oncotarget, 8/23: 38022–43.

Nam, J., Son, S., Park, K. S., Zou, W., Shea, L. D., & Moon, J. J. (2019). ‘Cancer nanomedicine for combination cancer immunotherapy’, Nature Reviews Materials, 4/6: 398–414. Nature Publishing Group.

Narayan, R. S., Molenaar, P., Teng, J., Cornelissen, F. M. G., Roelofs, I., Menezes, R., Dik, R., et al. (2020). ‘A cancer drug atlas enables synergistic targeting of independent drug vulnerabilities’, Nature Communications, 11/1: 2935. Nature Publishing Group.

Palmer, A. C., & Sorger, P. K. (2017). ‘Combination Cancer Therapy Can Confer Benefit via Patient-to-Patient Variability without Drug Additivity or Synergy’, Cell, 171/7: 1678–1691. e13.

Patel, S. A., & Minn, A. J. (2018). ‘Combination Cancer Therapy with Immune Checkpoint Blockade: Mechanisms and Strategies’, Immunity, 48/3: 417–33.

Subramanian, A., Tamayo, P., Mootha, V. K., Mukherjee, S., Ebert, B. L., Gillette, M. A., Paulovich, A., et al. (2005). ‘Gene set enrichment analysis: A knowledge-based approach for interpreting genome-wide expression profiles’, Proceedings of the National Academy of Sciences, 102/43: 15545–50. Proceedings of the National Academy of Sciences.

Sun, W., Sanderson, P., & Zheng, W. (2016). ‘Drug combination therapy increases successful drug repositioning’, Drug discovery today, 21/7: 1189–95.

Torkamannia, A., Omidi, Y., & Ferdousi, R. (2022). ‘A review of machine learning approaches for drug synergy prediction in cancer’, Briefings in Bioinformatics, bbac075.

Wu, L., Wen, Y., Leng, D., Zhang, Q., Dai, C., Wang, Z., Liu, Z., et al. (2022). ‘Machine learning methods, databases and tools for drug combination prediction’, Briefings in Bioinformatics, 23/1: bbab355.

Yang, W., Soares, J., Greninger, P., Edelman, E. J., Lightfoot, H., Forbes, S., Bindal, N., et al. (2013). ‘Genomics of Drug Sensitivity in Cancer (GDSC): a resource for therapeutic biomarker discovery in cancer cells’, Nucleic Acids Research, 41 /Database issue: D955–961.

Zheng, S., Aldahdooh, J., Shadbahr, T., Wang, Y., Aldahdooh, D., Bao, J., Wang, W., et al. (2021). ‘DrugComb update: a more comprehensive drug sensitivity data repository and analysis portal’, Nucleic Acids Research, 49/W1: W174–84.

## Supplementary References

Domanskyi, S., Hakansson, A., Bertus, T. J., Paternostro, G., & Piermarocchi, C. (2021). ‘Digital Cell Sorter (DCS): a cell type identification, anomaly detection, and Hopfield landscapes toolkit for single-cell transcriptomics’, PeerJ, 9: e10670.

Domanskyi, S., Szedlak, A., Hawkins, N. T., Wang, J., Paternostro, G., & Piermarocchi, C. (2019). ‘Polled Digital Cell Sorter (p-DCS): Automatic identification of hematological cell types from single cell RNA-sequencing clusters’, BMC bioinformatics, 20/1: 369.

